# Protein SUMOylation promotes cAMP-independent EPAC1 activation

**DOI:** 10.1101/2024.01.08.574738

**Authors:** Wenli Yang, Fang C. Mei, Wei Lin, Mark A. White, Li Li, Yue Li, Sheng Pan, Xiaodong Cheng

**Author notes:** Corresponding author: Xiaodong Cheng. **Email:**. These authors contributed equally (Wenli Yang, Fang C. Mei). Cell Therapy Manufacturing Center. 2130 W Holcombe Blvd, Houston TX, 77030.

## Abstract

Exchange protein directly activated by cAMP (EPAC1) mediates the intracellular functions of a critical stress-response second messenger, cAMP. Herein, we report that EPAC1 is a cellular substrate of protein SUMOylation, a prevalent stress-response posttranslational modification. Site-specific mapping of SUMOylation by mass spectrometer leads to identifying K561 as a primary SUMOylation site in EPAC1. Sequence and site-directed mutagenesis analyses reveal a functional SUMO-interacting motif required for cellular SUMOylation of EPAC1. SUMO modification of EPAC1 mediates its heat shock-induced Rap1/2 activation in a cAMP-independent manner. Structural modeling and molecular dynamics simulation studies demonstrate that SUMO substituent on K561 of EPAC1 promotes Rap1 interaction by increasing the buried surface area between the SUMOylated receptor and its effector. Our studies identify a functional SUMOylation site in EPAC1 and unveil a novel mechanism in which SUMOylation of EPAC1 leads to its autonomous activation. The findings of SUMOylation-mediated activation of EPAC1 not only provide new insights into our understanding of cellular regulation of EPAC1 but also will open up a new field of experimentation concerning the cross-talk between cAMP/EPAC1 signaling and protein SUMOylation, two major cellular stress response pathways, during cellular homeostasis.

## Introduction

Posttranslational modification with small ubiquitin-related modifier (SUMO) proteins is an essential and widespread regulatory mechanism for cellular homeostasis (1). A significant portion of the human proteome has been identified to undergo SUMO modifications (2). These SUMOylated proteins are involved in virtually all known cellular processes, including cell division, chromosome segregation, DNA replication/repair, gene transcription, nuclear transport, and signal transduction(1). SUMOylation regulates these cellular processes by altering the target protein’s activity, localization, stability, or interacting ability with binding partners (3). SUMOylation has long been associated with stress responses, integrating a diverse array of cellular stress signals that trigger rapid changes in global protein SUMOylation (4-7). Dysregulation of cellular SUMOylation has been implicated in the development of human diseases such as atherosclerosis, autoimmune diseases, cancer, diabetes, heart failure, and neurological disorders (8).

In parallel, the cAMP second messenger is a primary stress-response signal that plays essential roles in diverse biological functions under physiological and pathophysiological conditions. In multi-cellular organisms, the effects of cAMP are mainly transduced by two ubiquitously-expressed intracellular cAMP receptor families, the cAMP-dependent protein kinase (PKA) and the exchange protein directly activated by cAMP (EPAC) (9, 10). Between EPAC isoforms, EPAC1 and EPAC2, EPAC1 is ubiquitously expressed in all tissues, while EPAC2 has a limited tissue distribution, detected predominantly in the brain, pancreas and adrenal gland (11, 12). Despite acting on the same immediate down-stream effectors, the Ras superfamily small GTPases Rap1 and Rap2, cellular functions of EPAC1 and EPAC2 are frequently non-redundant due to their distinct tissue distribution and the ability to form discrete signalosomes at various cellular loci through interaction with specific cellular partners (13-16). Although protein SUMOylation and EPAC signaling were discovered around the same time more than two decades ago, crosstalk between cAMP/EPAC signaling and protein SUMOylation, two central cellular stress-response mechanisms, has not been explored extensively. Our recent study demonstrated that cAMP acts through EPAC1 to promote cellular SUMOylation via regulating the formation of biomolecular condensates enriched with SUMO processing enzymes and substrates, connecting the two major stress-response pathways (17). However, attempts to determine whether EPAC1 can undergo SUMO modification have not been successful. Consequently, the effects of EPAC1 SUMOylation remain unknown. In this study, we describe a novel finding that EPAC1 is a SUMO target protein and that SUMOylation of EPAC1 promotes its guanine exchange activity in a cAMP-independent manner.

## Results

### EPAC1 contains SUMO-consensus motifs (SCMs) and can be SUMOylated in cells

Many SUMO-modified proteins contain acceptor lysines within a ΨKX[D/E] SCM or an inverted ([E/D]XKΨ) SCM, where Ψ is a large hydrophobic amino acid, and X is any amino acid (18). When the human EPAC1b sequence was analyzed using a SUMOylation prediction algorithm (19), several putative SCMs, as well as a SUMO-interacting motif (SIM) (20, 21), were identified (**Table S1**). To test if EPAC1 is SUMOylated in cells, we subjected cells to heat shock treatment, known to induce robust cellular SUMOylation (6). We subsequently probed the status of EPAC1, endogenously in Human Umbilical Vein Endothelial Cells (HUVECs) or ectopically expressed in HEK293 cells, which expresses a very low level of EPAC1 (9), by immunoblotting using a monoclonal EPAC1 antibody, 5D3. Incubation of HUVECs or HEK293/EPAC1-Flag cells at 43 °C led to time-dependent accumulation of higher molecular weight EPAC1 immunoblotting signals above the ∼100 Kd EPAC1 native protein band (**Fig. 1A, B**). Similar results were obtained in HeLa cells transfected with an EPAC1-Flag/HA containing pOZ vector (22) (**Supplementary Fig. 1A)**. Importantly, when cell lysates were treated with Sentrin-specific protease 1 (SENP1), a deSUMOylation protease, the higher molecular weight EPAC1 immunoblotting signals were markedly reduced (**Fig. 1C, Supplementary Fig. 1B**), suggesting that some of these higher molecular weight EPAC1 post-translational modification (PTM) signals are indeed originated from SUMOylated EPAC1.

**Figure 1.**
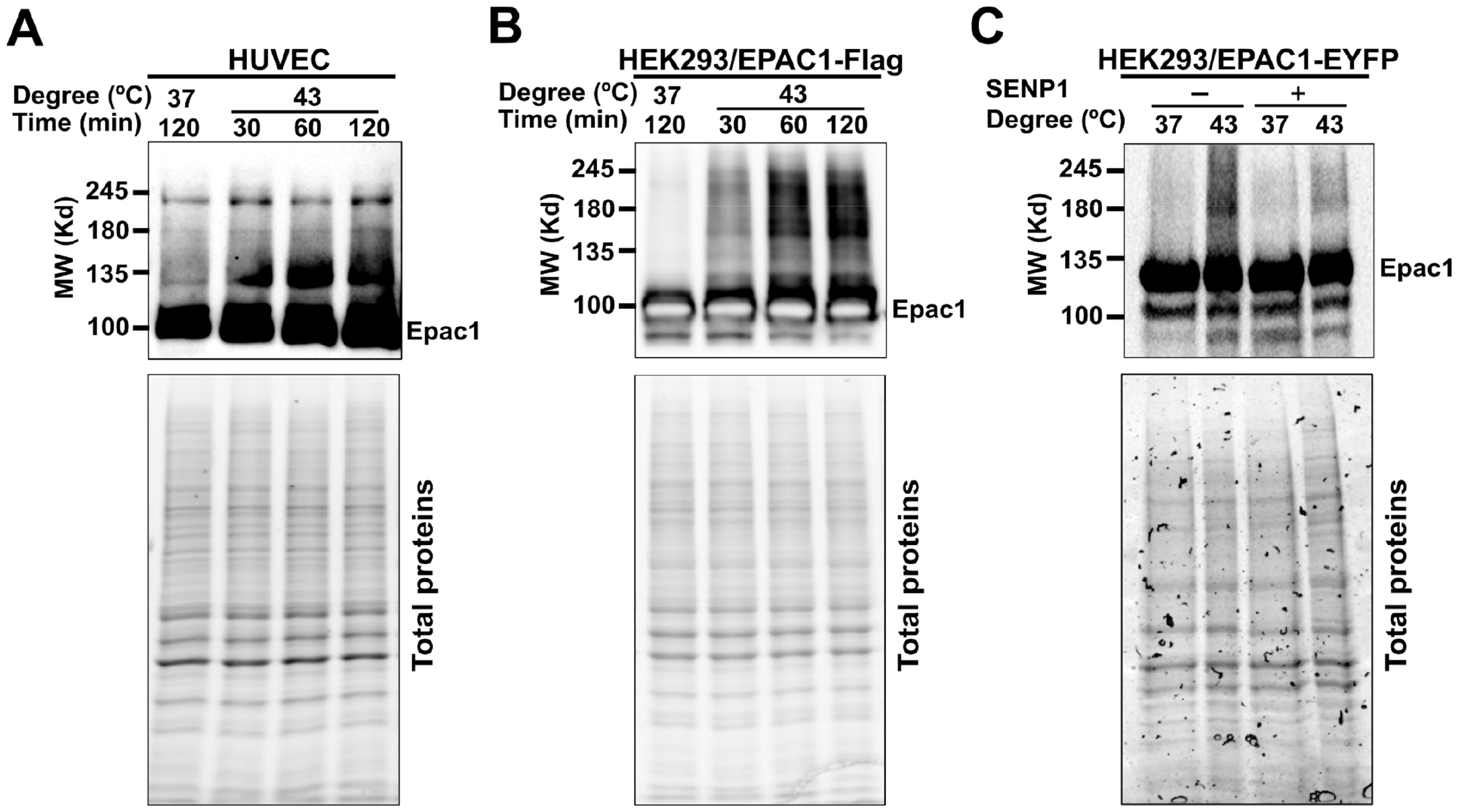
Heat shock promotes EPAC1 post-translational modifications (PTM). Levels of cellular EPAC1 PTM probed by immunoblotting using anti-EPAC1 antibody in HUVEC (A) and HEK293/EPAC1-Flag (B) in response to heat shock as a function of time. (C) Levels of cellular EPAC1 PTM in HEK293/EPAC1-EYFP cells probed by immunoblotting using anti-EPAC1 antibody, with and without heat shock (30 min) and with or without SENP1 (220 nM) treatment at 37 °C for 20 min. Similar results were obtained from at least three independent experiments.

### Identification of an EPAC1 SUMOylation site by mass spectrometry (MS)

Encouraged by the finding of heat shock-induced, SENP1-sensitive EPAC1 PTMs, we mapped potential SUMOylation sites in EPAC1. Site-specific identification of SUMOylation by MS remains technically challenging owing to the dynamic nature of cellular SUMOylation (23) and the lack of naturally occurring tryptic sites near the C-terminal tail of SUMO proteins as in the case of ubiquitin (2). To overcome these challenges, we mutated the Q89 residue of SUMO3 to K so that tryptic digestion of conjugated SUMO3(Q89K) would leave a “signature peptide” containing a TGG adduct attached to the SUMOylated lysine on the substrate, allowing robust and unambiguous identification of the SUMOylation site using MS. We ectopically expressed the EPAC1-His_10_ and SUMO3(Q89K) in Hela cells simultaneously using a biscistronic lentivector and performed denaturing His-tag pull-down using lysate from cells treated with or without 30 min 43 °C heat shock (**Fig. 2A**). Immunoblotting analysis of the pull-down samples by an anti-SUMO2/3 antibody revealed that heat shock treatment led to a robust increase in SUMO2/3-containing high molecular weight bands (**Fig. 2B**). These results further support that EPAC1 can be SUMOylated, and heat shock promotes EPAC1 SUMOylation. MS analysis of the His-tag pull-down eluent of the heat shock sample led to the unambiguous identification of the K561 as a SUMOylation site in EPAC1 with a less than 1% FDR (**Fig. 2C**). The overall MS data quality was excellent with a sequence coverage for EPAC1 more than 90%. The MS spectra for peptides spanning the K561 were all high quality, both for modified and unmodified peptides. This allowed us to estimate the stoichiometry of K561 SUMOylation at ∼9% based on the PSM (peptide-spectrum match) counts.

**Figure 2.**
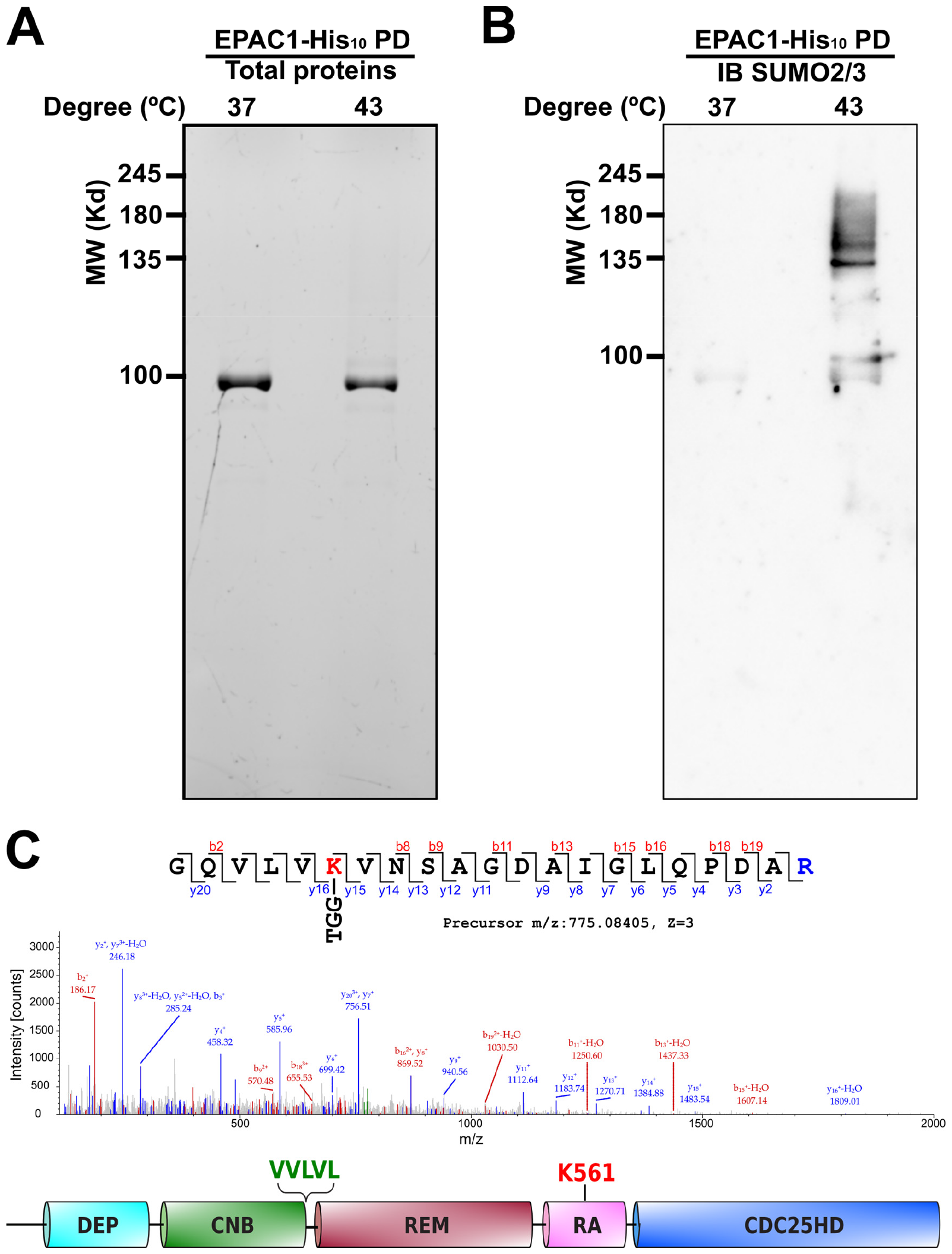
EPAC1 is a SUMO target protein. Protein gel (A) and anti-SUMO2/3 immunoblotting (B) of nickel affinity pull-down of HEK293/pCDH-CMV-EPAC1-His_10_-IRES-SUMO3(Q89K) lysates at 37 and 43 °C. (C) MS/MS spectrum and corresponding peptide sequence of EPAC1 SUMOylation site K561 (red). The domain structure of EPAC1 with a predicted SUMO-interacting motif highlighted in green and K561 highlighted in red.

### An EPAC1 SUMO interacting motif (SIM), but not cAMP, is required for SUMO modification

Many SUMOylation substrates also contain SIM that interacts with free or conjugated SUMO (20, 21). In certain SUMOylation targets, the SUMO-binding property of the SIM contributes to substrate recognition and is critical for SUMOylation (24, 25). Since EPAC1 has a well-defined SIM, ^320^VVLVL^324^, we hypothesized that the EPAC1 SIM might play a role in EPAC1 SUMOylation. We replaced the EPAC1 SIM motif, VVLVL, with alanines to generate the EPAC1(SIM/5A) mutant to test this hypothesis. We ectopically expressed WT EPAC1-APEX2 and EPAC1(SIM/5A)-APEX2 in HEK293 cells and performed affinity pull-down using anti-SUMO2/3 antibody. Immunoblotting analysis of the pull-down samples by an anti-EPAC1 antibody showed that mutation of the EPAC1 SIM led to the complete abolishment of high molecular SUMOylated EPAC1 bands seen in WT EPAC1-APEX2 fusion under both normal and heat shock conditions (**Fig. 3A**). These results suggest that EPAC1 SIM is essential for EPAC1 SUMOylation.

**Figure 3.**
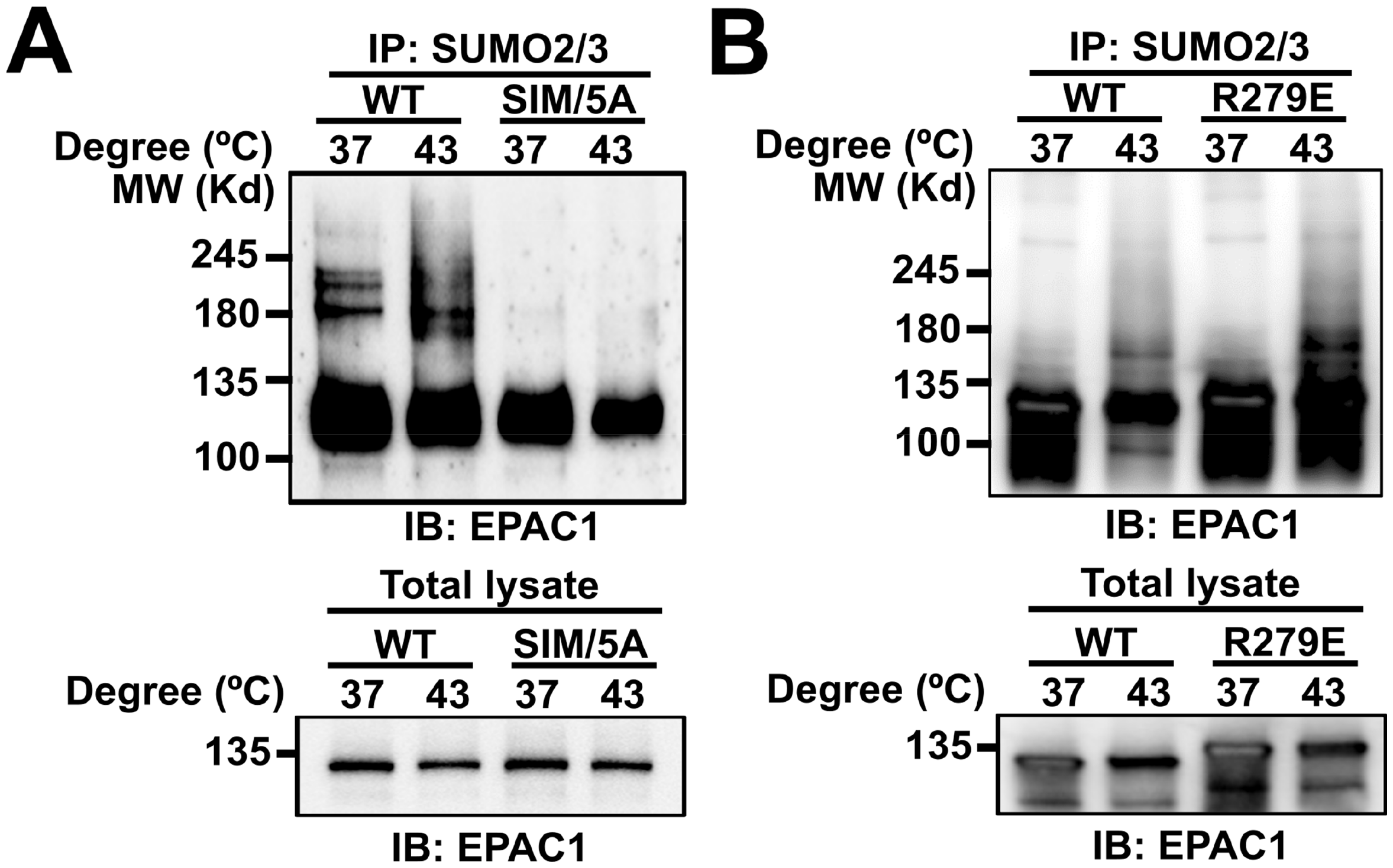
SIM-dependent and cAMP-independent SUMOylation of EPAC1. (A) Anti-EPAC1 immunoblotting of SUMO2/3 affinity pull-down of HEK293/EPAC1-APEX2 and HEK293/EPAC1(SIM/5A)-APEX2 after incubation at 37 or 43 °C for 30 min. (B) Anti-EPAC1 immunoblotting of SUMO2/3 affinity pull-down of HEK293/EPAC1-EYFP and HEK293/EPAC1(R279E)-EYFP after incubation at 37 or 43 °C for 30 min.

Since EPAC1 SIM overlaps with a previously identified conserved switchboard sequence required for the autoinhibition of EPAC1 (26), we asked if EPAC1 activation by cAMP binding is necessary for heat-shock-induced EPAC1 SUMOylation. To address this question, we used a well-characterized EPAC1 mutant, EPAC1(R279E), with a single point mutation of a conserved and critical residue, Arg279, that is required for cAMP-binding (27). Anti-SUMO2/3 affinity purification followed by EPAC1 immunoblotting showed that unlike EPAC1(SIM/5A), EPAC1(R279E) mutant retained the capability of being SUMOylated in response to heat shock as WT EPAC1 (**Fig. 3B**). These results suggest that heat shock-mediated EPAC1 SUMOylation does not require cAMP binding.

### Biomolecular condensate, not RanBP2, is involved in heat shock-induced EPAC1 SUMOylation

EPAC1 interacts with RanBP2 (28, 29), a bona fide SUMO E3 ligase (30). The close association between EPAC1 and RanBP2 raises the possibility that RanBP2 is involved in EPAC1 SUMOylation. However, when we performed the EPAC1 affinity pull-down experiment and probed the RanBP2, we found that heat shock led to a significant reduction of EPAC1 and RanBP2 interaction (**Supplementary Fig. 2**). This result suggests that RanBP2 is unlikely an E3 ligase that catalyzes the SUMOylation of EPAC1 in response to heat shock. On the other hand, we ectopically expressed EPAC1-EYFP and mRuby tagged ubiquitin-like modifier activating enzyme 2 (UBA2), the catalytic subunit of the SUMO E1 enzyme, in HEK293 cells and performed confocal imaging. At 37 °C, EPAC1-EYFP signals were mostly diffused. A few puncta were present in the cytosol but mostly absent in the nuclear compartment, while dispersed UBA2 speckles were observed in the nuclei. In response to heat shock stimulation, mRuby-UBA2 coalesced to form larger nuclear condensates. During this process, we also observed an increased formation of EPAC1-EYFP puncta, particularly in the nuclei that were superimposable with the mRuby-UBA2 in a time-dependent manner (**Fig. 4**). These results, coupled with our previous findings that EPAC1 activation enhances cellular SUMOylation via promoting the formation of biomolecular condensates enriched with SUMOylation machinery (17), suggest that in addition to induce SUMOylation promoting condensates, EPAC1 itself is a SUMO substrate during the process. To further affirm the importance of nuclear condensate forming capability for heat-shock-induced EPAC1 SUMOylation, we tested the behavior of EPAC1(R279E)-EYFP and EPAC1(SIM/5A)-EYFP in response to heat shock. Our previous study shows that R279E mutation abolishes cAMP-induced EPAC1 nuclear condensates formation (17). However, EPAC1(R279E)-EYFP, similar to WT EPAC1-EYFP, formed robust nuclear condensates that were superimposable with the mRuby-UBA2 condensates in response to heat shock. On the other hand, EPAC1(SIM/5A)-EYFP is mainly cytosolic, and mutation of the SIM abolished EPAC1’s ability to form nuclear condensates in response to heat shock (**Supplementary Fig. 3)**. The nuclear condensate formation capability of these mutants matched their SUMOylation status in response to heat shock, as shown in **Figure 3**, further supporting that nuclear condensates were involved in heat-shock-induced EPAC1 SUMOylation.

**Figure 4.**
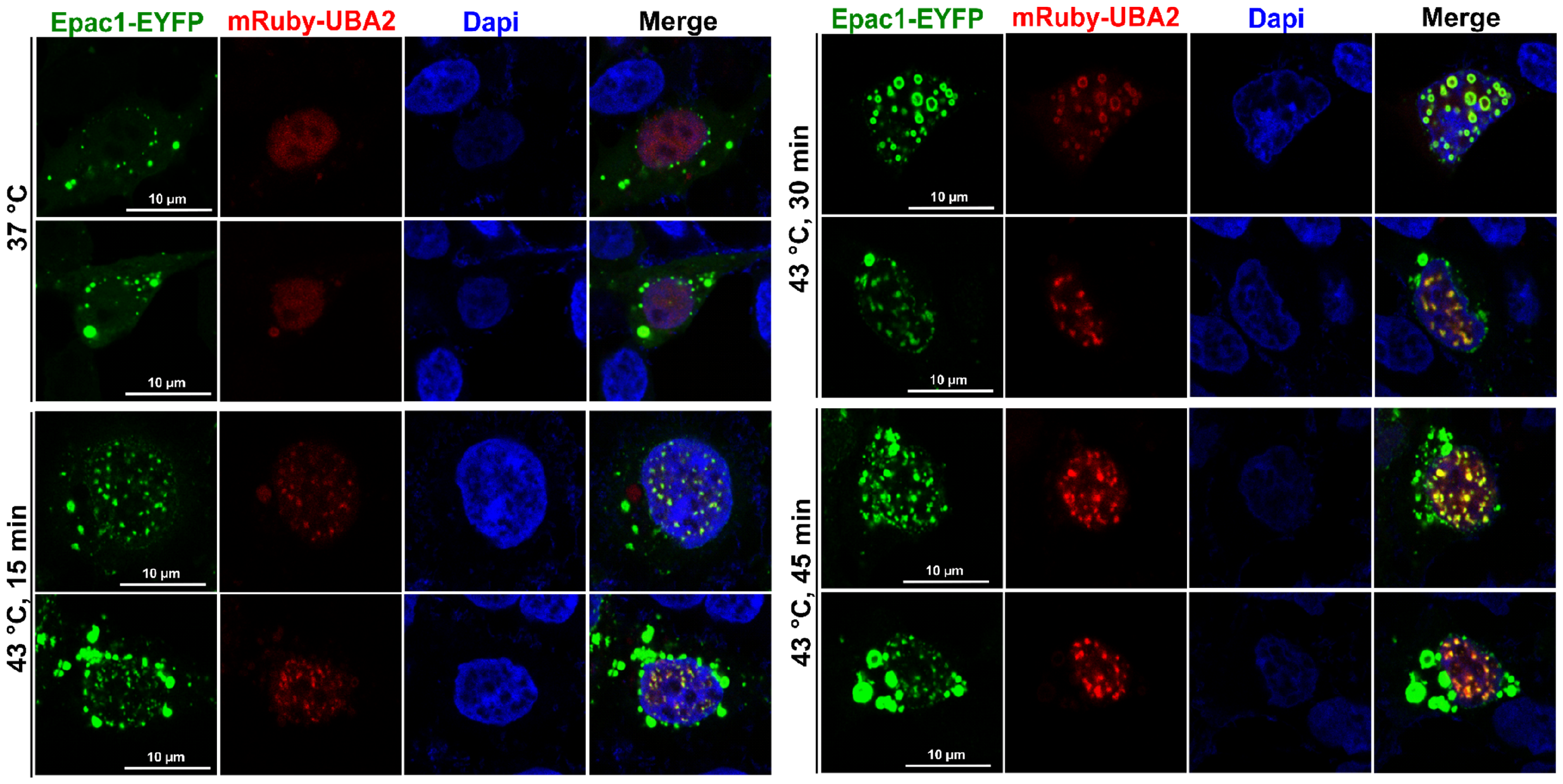
Heat shock-induced formation of nuclear EPAC1-EYFP/mRuby-UBA2 condensates. Confocal images of HEK293 cells ectopically expressing EPAC1-EYFP and mBuby-UBA2 in response to heat shock treatment as a function of time.

### EPAC1 K561 SUMOylation is responsible for heat-shocked induced Rap GTPase activation

Protein SUMOylation is known to modulate the activity, stability, cellular localization, or interacting ability of the target protein (3). To ascertain the functionality of EPAC1 K561 SUMOylation, we ask if K561 SUMOylation affects EPAC1’s guanine nucleotide exchange activity. Since under normal conditions, only a tiny fraction of EPAC1 is SUMOylated, we first tested if heat shock treatment, known to induce robust cellular SUMOylation, affected Rap1/2 activation. **Figure 5A** shows heat shock-induced significant increases in both Rap1-GTP and Rap2-GTP levels. To test if EPAC1 is responsible for the heat-shocked mediated Rap GTPase activation, we silenced EPAC1 expression via RNAi and compared heat shock-induced Rap activation in HUVECs cells treated with control or EPAC1-specific RNAi. Knocking down EPAC1 blocked heat-shocked mediated Rap GTPase activation (**Fig. 5B**), suggesting that EPAC1 is a direct downstream effector of heat shock in terms of Rap activation.

**Figure 5.**
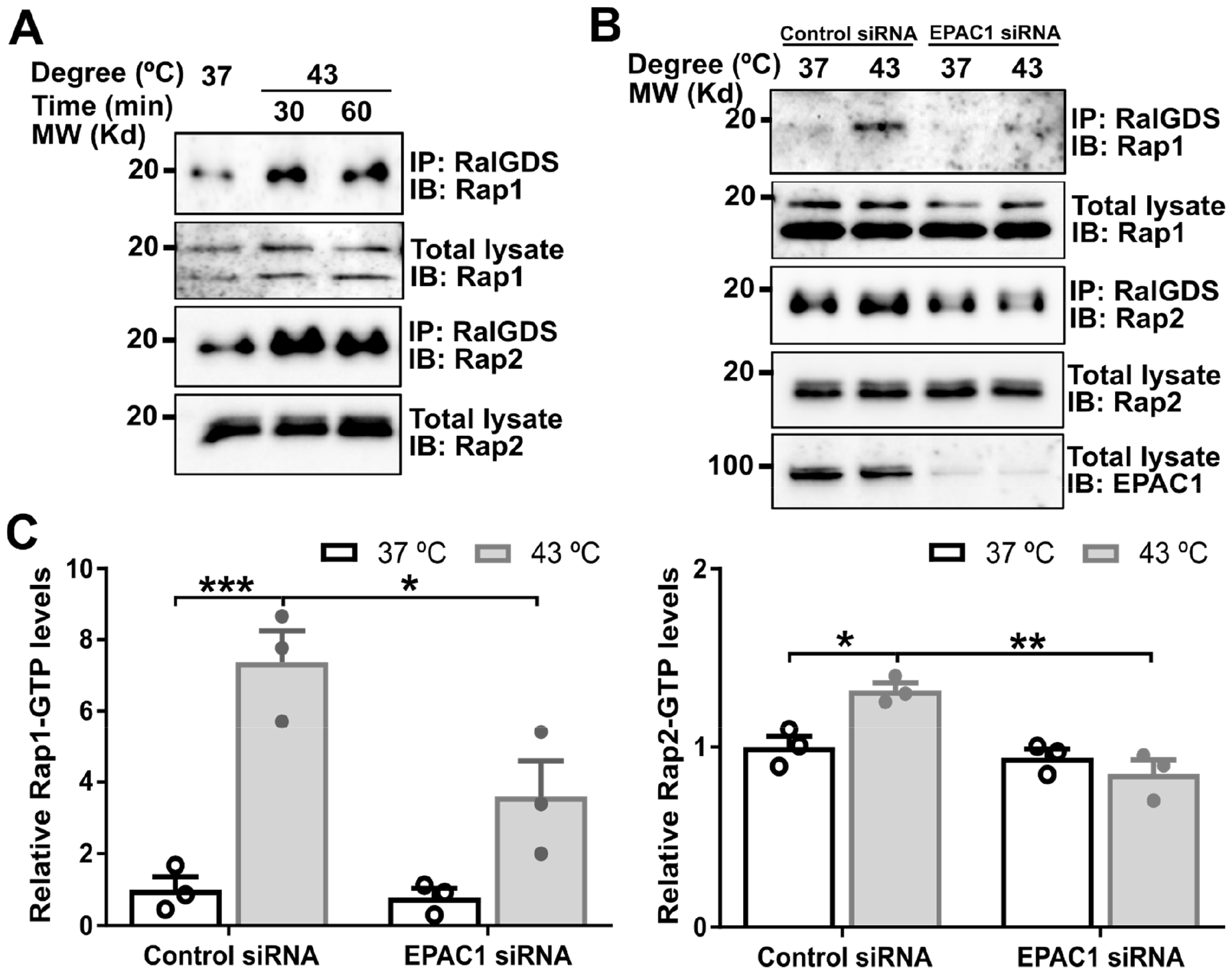
EPAC1 is responsible for heat shock-induced activation of Rap small GTPases in HUVEC. (A) heat shock promoted cellular activation of Rap1 and Rap2. (B) Silencing of EPAC1 abolished heat shock-mediated Rap activation. (C). Quantification of cellular Rap1-GTP and Rap2-GTP levels in C. Data are presented as Mean ± SEM (N = 3).

To test if heat shock activates EPAC1 via increases in intracellular cAMP levels, we determined the intracellular cAMP concentration in HUVEC and HEK293 cells in response to heat shock. Our data showed that heat shock did not significantly affect the intracellular cAMP levels in both cells (**Supplementary Fig. 4**). These findings led us to ask if EAPC1 K561 SUMOylation contributed to heat shock-induced Rap activation. When wild-type EPAC1-EYFP or an EPAC1(K561R)-YFP mutant was ectopically expressed in HEK293 cells, heat shock was able to promote Rap activation in HEK293/EPAC1-EYFP cells. However, this effect was abolished entirely in HEK293/EPAC1(K561R)-YFP cells (**Fig. 6A**), suggesting that EPAC1 K561 SUMOylation is responsible for heat-shocked induced Rap GTPase activation.

**Figure 6.**
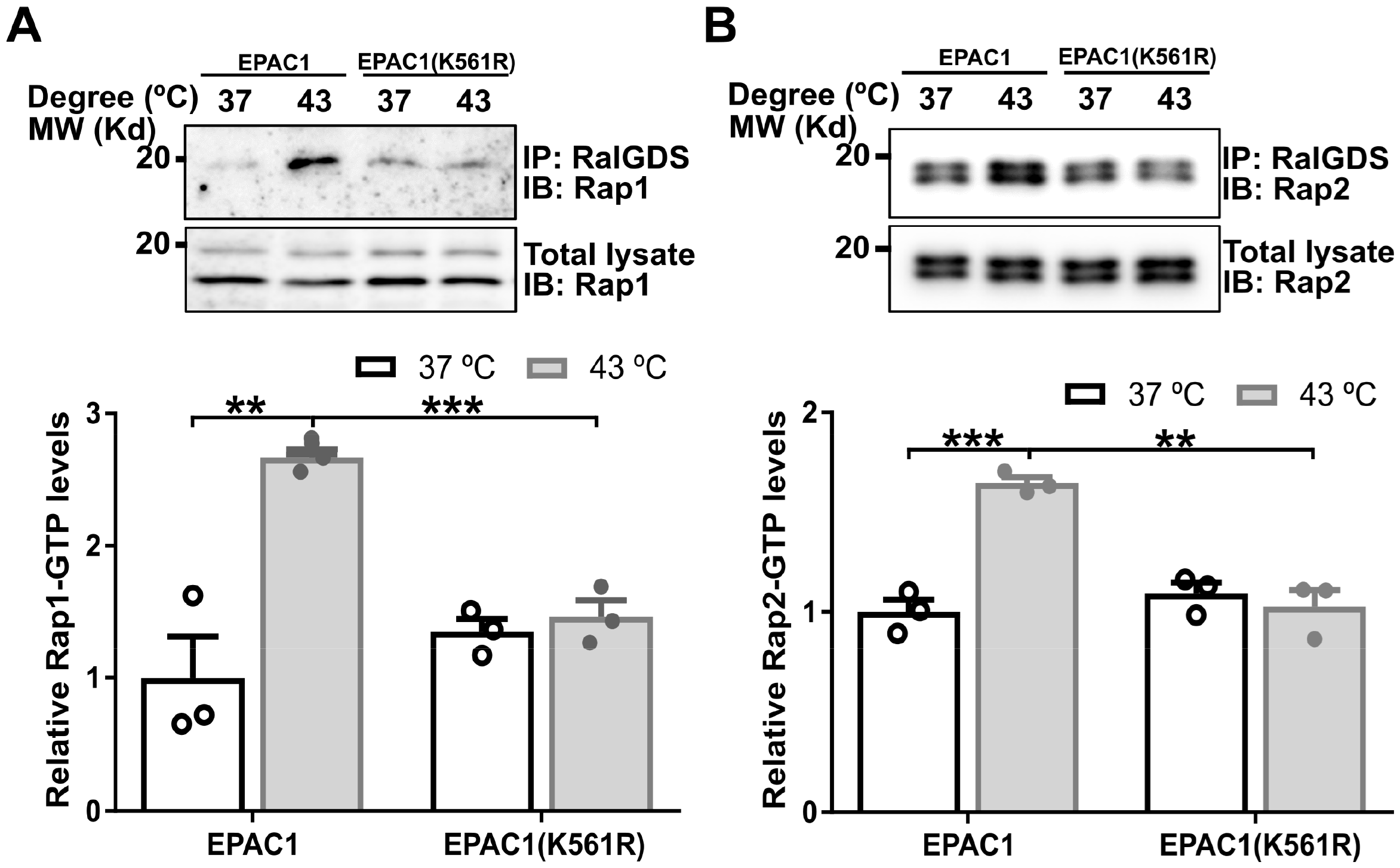
K561 SUMOylation of EPAC1 is essential for heat shock-induced activation of Rap small GTPases. (A) K561R mutation abolished heat shock-promoted cellular activation of Rap1 (A) and Rap2 (B). Data are presented as Mean ± SEM (N = 3).

### K561 SUMOylation of EPAC1 enhances Rap GTPase binding

To understand the structural basis of EPAC1(K561) SUMOylation mediated Rap GTPase activation, we performed structural modeling and molecular dynamics (MD) simulation analyses to generate K561 SUMOylated EPAC1 structures in apo conformation (EPAC1(K561)-SUMO) or active conformation with the Rap1 effector bound (EPAC1(K561)-SUMO:cAMP:Rap1). Due to the presence of Rap1 in the active conformation, the possible position of the K561-SUMO moiety is restricted, while the initial position of the SUMO moiety is not restrained by its covalent bond to EPAC1 K561 in the apo conformation (**Supplementary Fig. 5A**). Despite this difference, the positions of the SUMO moiety of the two models remain similar after MD simulation with the SUMO orientation in EPAC1(K561)-SUMO shifted slightly in the absence of the Rap1 but remained in roughly the same position as in EPAC1(K561)-SUMO:cAMP:Rap1 (**Supplementary Fig. 5B & C**). The SUMO moiety seemed to position itself away from the N-terminal regulatory lobe (**Supplementary Fig. 6A**) and the Rap1 binding site in the CDC25HD domain, leaving it accessible for effector binding (**Supplementary Fig. 5D**). These results suggest that the SUMO moiety likely assumes a stable orientation before Rap1 binding. Analysis of the EPAC1(K561)-SUMO:cAMP:Rap1 revealed that the SUMO moiety moved slightly toward the bound Rap1 to make direct contact (**Supplementary Fig. 5D**). At the same time, the Rap1 rotated somewhat to accommodate the SUMO but remained securely bound to the EPAC1 CDC25HD domain (**Supplementary Fig. 6B**). In addition to the added interaction between SUMO and Rap1, the (K561)-SUMO substituent also increased the surface area between Rap1 and EPAC1 CDC25HD domain. As a consequence, the Rap1 footprint on the EPAC1(K561)-SUMO covered a significantly larger surface area of 2,227 Å^2^ (**Fig. 7A & B**) as compared to a buried surface area of 1,486 Å^2^ between Rap1 and EPAC1 alone (**Fig. 7C & D**).

**Figure 7.**
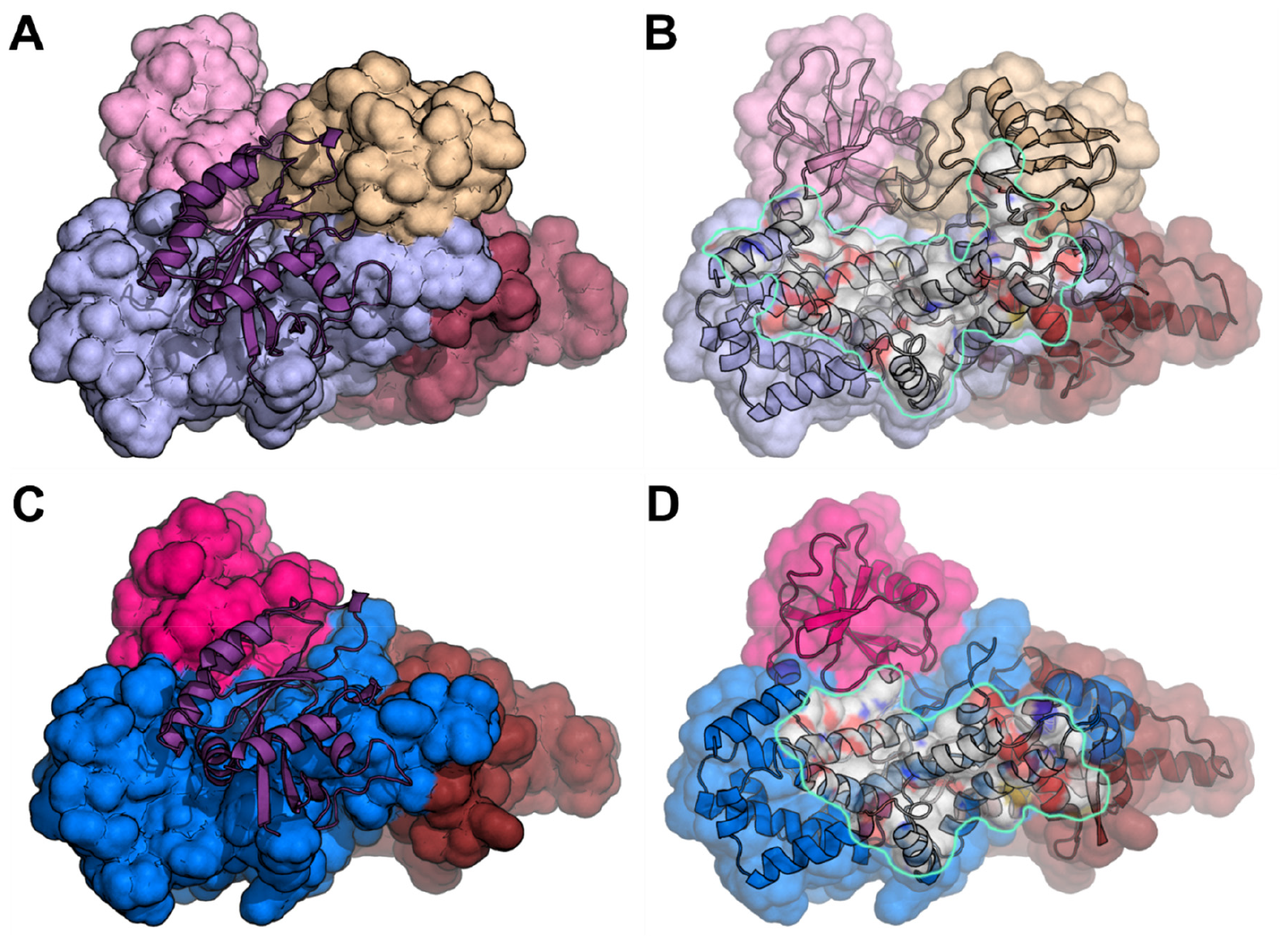
K561 SUMOylation of EPAC1 promotes Rap1 interaction. (A) The MD EPAC1^CT^-SUMO:Rap1 model after 30 ns of Molecular Dynamics showing the Rap1 (cartoon) bound to EPAC1^CT^-SUMO. (B) The same view as in A with Rap1 removed, and the Rap1 interface highlighted. The Rap1 total buried surface area is 2,227 Å^2^. (C) Corresponding view of the full-length EPAC1:cAMP:Rap1 model’s catalytic domains after 30 ns of Molecular Dynamics showing the Rap1 (cartoon) bound to EPAC1 (For clarity, the regulatory domains are not shown). (D) The same view as in C with Rap1 removed, and the Rap1 interface highlighted. The Rap1 total buried surface area is 1,486 Å^2^. The EPAC1 domains are colored as previously (REM: maroon; RA: pink; CDC25HD: blue). The SUMO is beige, and the Rap1 is purple.

## Discussion

The universal second messenger, cAMP is an ancient stress response signal conserved from bacterium to human and vital for optimal adaptation. Conversely, protein SUMOylation is a prevalent posttranslational modification essential for maintaining cellular homeostasis (31). Recent studies demonstrate that cAMP, acting through EPAC1, can modulate cellular SUMOylation by promoting the formation of biomolecular condensate enriched with components of SUMOylation machinery (17, 32). These findings establish a crucial nexus linking two central cellular stress-response mechanisms. In this study, we reveal that SUMOylation of EPAC1 on K561 of the Ras association (RA) domain enhances EPAC1 catalytic activity by providing an additional binding surface for its effector small GTPase Rap1/2. The unequivocal identification of a functional SUMOylation site in EPAC1 offers another mechanism connecting protein SUMOylation and cAMP signaling. It is worth noting that previous studies have reported that EPAC1 interacts with RanBP2 (25, 26), a bona fide SUMO E3 ligase (27). However, attempts to show SUMOylation of EPAC1 were not successful (26). This is most likely because the level of cellular EPAC1 SUMOylation is very low and highly dynamic. To overcome this challenge, we combined ectopic expressing a mutant SUMO3(Q89K) tag with heat shock to promote cellular SUMOylation. These modifications allow us to identify K561 as a primary SUMOylation site in EPAC1.

Lysine 561 is located within the RA domain, which displays significant sequence diversity between the two isoforms (33). RA domain folds into a ubiquitin alpha/beta roll superfold (34) and has been found in various proteins to function mainly as a protein interaction scaffold (35). Previous studies have demonstrated that the RA domain contributes to isoform-specific functions of EPACs. RA domain in EPAC2 interacts with RAS and mediates EPAC2 translocation to the plasma membrane and activation (36-38). A G706R rare coding mutation in the EPAC2 RA domain has been found in several autistic patients (39). Functional analyses have revealed that G706R mutation impairs RAS interaction and selectively reduces basal dendrite complexity in cortical pyramidal neurons (40). Conversely, the RA domain of EPAC1 is known to interact with β-arrestin2 and differentially regulates cardiac hypertrophic signaling mediated by β-adrenergic receptor subtypes (41). In addition, EPAC1 RA has also been shown to mediate the interaction with Ran-GTP and RanBP2 proteins and to target EPAC1 to the nuclear membrane (28). However, a subsequent study disputes the conclusions (29). Our finding that SUMO modification of K561 in the EPAC1 RA domain enhances the interaction and activation of Rap GTPases further expands the role of RA domains in isoform-specific functions of EPACs. It will be interesting to test the effects of K561 SUMOylation on EPAC1’s interaction with β-arrestin2 or RanBP2.

MD analyses of the EPAC1(K561)-SUMO revealed that the K561-SUMO moiety assumed a stable conformation without the effector, Rap1, despite without steric restriction. In the presence of Rap1, the SUMO moiety moved slightly to make direct contact with the bound Rap1, which provided an extra binding interface to promote Rap1 binding to EPAC1(K561)-SUMO as compared to unmodified EPAC1. The measured Rap1 buried SASA increased from 1,486 Å^2^ to 2,227 Å^2^ with SUMOylation. Structural analysis of SUMOylated proteins remains challenging due to lacking a general approach to generating pure proteins with site-specific SUMOylation. Our current study demonstrates that MD analysis is effective in interrogating EPAC1 SUMOylation to provide structural insights into SUMO modification and may present a general structural methodology for the analysis of protein SUMOylation.

In addition to being a SUMOylation substrate, EPAC1 also contains a well-defined SUMO-interacting motif. Interestingly, EPAC1 SIM, ^320^VVLVL^324^, overlaps with the “switchboard” (SB) sequence ^321^VLVLE^325^ that is reported to be critical for maintaining the proper orientation between the regulatory and catalytic halves of EPAC1 (26). Activation of EPAC1 by cAMP leads to a hinge motion that allows the translocation of the SB to become the “lid” of the cAMP binding pocket (42, 43). Surprisingly, our study reveals that EPAC1 SIM is necessary for EPAC1 SUMO modification, suggesting that it also acts as a switchboard for EPAC1 SUMOylation. The discovery that EPAC1 can be SUMOylated in cells and the identification of K561 as a primary EPAC1 SUMO modification site, coupled with the findings that EPAC1 contains a functional SIM, places EPAC1 among many known SUMO-interacting proteins that are themselves covalently SUMOylated (44). The simultaneous presence of SUMO conjugation and SIM in single proteins may contribute to liquid-liquid phase separation (LLPS) behind the formation of a family of membraneless biomolecular condensates to regulate cellular processes spatially and temporally via coordinating the assembly of large protein complexes/networks (31). Considering the recent findings that EPAC1 can undergo LLPS to form cellular condensates involved in the regulation of protein SUMOylation (17) and histone transcription (45), it will be essential to test if EPAC1 SUMOylation is involved in the formation or regulation of EPAC1 condensates. Regardless, results from this study establish a functional role of protein SUMOylation in EPAC1-mediated signaling.

## Materials and Methods

### Reagents

Anti-EPAC1 antibody 5D3 (Cell Signaling Technology, #4155), anti-SUMO2/3 antibody (MBL Life Science, Catalog no. M114-3), anti-Ran BP2 antibody (Santa Cruz Biotechnology, Inc., Catalog no. sc-74518), Anti-Rap1 antibody (Santa Cruz Biotechnology, Inc., Catalog no. sc-65) and Anti-Rap2 antibody (BD Biosciences, Cat# 610215) were used in this study. Dulbecco’s modified Eagle’s medium (DMEM) high glucose (Catalog no. D5796), fetal bovine serum (FBS) (Catalog no. F2442), N-ethylmaleimide (NEM) (Catalog no. E3876), Ni Sepharose Fast Flow (Catalog no. GE17-5318), poly-l-lysine solution (0.01%, catalog no. P4707) and FluorSave Reagent (catalog no. 345789) were from MilliporeSigma. Antibiotic-antimycotic (100×) (Catalog no. 15240096), 4′,6-diamidino-2-phenylindole (DAPI) (Catalog no. 62248), and Lipofectamine 2000 (Catalog no. 11668-019) were from Thermo Fisher Scientific. cOmplete, Mini, EDTA-free Protease Inhibitor Cocktail Tablet was from Roche (catalog no. 11836170001). Protein A/G PLUS-Agarose (Catalog no. sc-2003) was from Santa Cruz Biotechnology Inc. Tris(2-carboxyethyl)phosphine hydrochloride (TCEP) (Catalog no. HR2-651) was from Hampton Research. Recombinant Human His6-SENP1 Catalytic Domain (SENP1, Catalog no. E-700) was from R&D Systems.

### Constructs

Human EPAC1b protein C-terminally tagged with a Flag or EYFP was described previously (46). pOZ-FH-C-EPAC1 construct encoding a C-terminal HA/Flag double epitope-tagged EPAC protein was previously described (22). EPAC1-V5-APEX2 construct was generated by inserting the EPAC1 gene into the pCDNA3-V5-APEX2 vector (47) using NotI and NheI restriction enzyme sites. The EPAC1-V5-APEX2 fragment was linearized with EcoRI and NotI and cloned into the pCDH-CMV-MCS-EF1α-Puro (System Biosciences, Catalog no. CD510B-1) lentiviral vector. A bicistronic construct expressing human EPAC1b with a C-terminal His_10_ tag and SUMO3(Q89K) was constructed using the pIRES2-EGFP vector (Clontech Catalog no. 632435). The EPAC1-His_10_-IRES-SUMO3(Q89K) fragment was linearized with EcoRI and NotI and cloned into the pCDH-CMV-MCS-EF1α-Puro lentiviral vector to generate the pCDH-CMV-EPAC1-His_10_-IRES-SUMO3(Q89K) plasmid.

### SUMOylation site and SUMO-interacting motif prediction

Putative SUMOylation sites and SUMO-interacting motifs of human EPAC1b were analyzed using a web-based Joined Advanced SUMOylation Site and SIM Analyser (http://www.jassa.fr/) (19). Candidates with “High cut-off” predictive scores or “DB hit” that matches a previously validated SUMOylation site, or SIM were selected as putative hits.

### Cell culture and transfection

HEK-293 (ATCC, Catalog no. CRL-1573) and HeLa (ATCC, Catalog no. CCL-2) cells were grown in DMEM supplemented with 10% heat-inactivated FBS (Invitrogen) at 37 °C, 5% CO_2_. HUVECs (Lonza, catalog no. C2519A) were maintained and subcultured in EGM-2 Endothelial Cell Growth Medium (Lanzo, catalog no. CC-3162) at 37°C in a 5% CO_2_ humidified incubator. Cell passages between 2 and 8 were used for experiments described in this study. For experiments involving RNA interference (RNAi), HUVECs at 70% confluence were transfected with EPAC1-specific (Thermo Fisher Scientific, Catalog no. 1299001) or nontargeting control Stealth RNAi siRNA oligonucleotides (Thermo Fisher Scientific, Catalog no. 12935300) at a final concentration of 50 nM. Cell transfection was performed using Lipofectamine 2000 according to the manufacturer’s instructions.

### Heat shock-induced EPAC1 posttranslational modification (PTM)

Near-confluent HUVEC, HeLa/pOZ-FH-C-EPAC1 (22), and HEK293 cells ectopically expressing EPAC1-Flag or EPAC1-EYFP grown in 12-well plates were subjected to heat shock treatment at 43 °C for various periods. After treatment, cells were washed twice with warm Dulbecco’s phosphate-buffered saline (DPBS) and lysed with 100 μl of 1× SDS sample buffer [50 mM tris (pH 6.8), 2% SDS, 0.1% bromophenol blue, 3% 2-ME, and 10% glycerol] with protease inhibitors and 20 mM NEM. Total cell lysates were collected and sonicated on ice using 15-W power output for three to four cycles of 5 seconds, with 5-s rests in between until entirely soluble. After heat denaturation at 95°C for 5 min, the samples were subjected to immunoblotting analysis using an anti-EPAC1 antibody. To test if EPAC1 PTM was sensitive to SUMO-specific deconjugating enzyme, heat shock treated cells were lysed with 1× cell lysis buffer without 20 mM NEM and then incubated with 220 nM recombinant Human Sentrin-specific protease 1 (SENP1) catalytic domain (R&D Systems, Catalog no. E-700) at 37 °C for 20 min before immunoblotting analysis.

### Nickel affinity pull-down of HEK293/pCDH-CMV-EPAC1-His_10_-IRES-SUMO3(Q89K) treated by HS

HEK-293 cells stably expressing EPAC1-His_10_ and SUMO3(Q89K) was generated using pCDH-CMV-EPAC1-His_10_-IRES-SUMO3(Q89K) lentiviral vector and used for the *in vivo* identification of the EPAC1 SUMOylation site via nickel affinity purification under denaturing conditions using a well-established procedure as described previously (48). Briefly, when eight 10 cm plates of HEK-293 cells stably expressing EPAC1-His_10_ and SUMO3(Q89K) reached 70%-80% confluence, four plates were subject to heat shock treatment at 43 °C for 30 min while the remaining four were kept at 37 °C. The cells were rinsed in PBS three times and lysed with 1 ml freshly prepared lysis buffer [10 mM Tris, 100 mM Sodium phosphate, pH 7.8, 400 mM NaCl, 6 M guanidine chloride, 20 mM imidazole, 0.5 mM EDTA, 0.5 mM PMSF, 10 mM NEM] directly on the plate. After gently rocking for 10 min, cell lysate was scrapped off, pooled, sonicated for 5 s, three times, on low power, and centrifuged at 16,000 g for 15 min to remove cell debris. The resultant supernatant was mixed with 45 μl of Sephorase 6 fast-flow nickel beads (GE 17-5318-02) equilibrated in the lysis buffer and incubated at room temperature with gentle mixing for three hours. The supernatant was carefully removed after centrifugation at 1,000 g for 3 min. The remaining beads were washed twice with lysis buffer containing 0.1% TritonX-100, twice with freshly prepared pH 8.0 wash buffer [10 mM Tris, 100 mM sodium phosphate, pH 8.0, 8 M Urea, 0.1% (vol/vol) Triton X-100, 5 mM β-mercaptoethanol], and three times with freshly prepared pH 6.3 wash buffer [10 mM Tris, 100 mM sodium phosphate, pH 6.3, 8 M Urea, 0.1% (vol/vol) Triton X-100, 5 mM β-mercaptoethanol]. EPAC1-His_10_ was eluted from the beads with elution buffer [150 mM Tris-HCl pH 6.7, 300 mM Imidazole, 5% (wt/vol) SDS, 30% (vol/vol) glycerol, 5 mM β-mercaptoethanol]. The eluents were loaded onto 10% SDS–polyacrylamide gel electrophoresis (SDS-PAGE) gel. Shortly after all the eluents were migrated into the gels, gel bands (1 to 2 cm) containing the total protein loading were excised after staining with Coomassie Blue. The gel bands were subjected to in-gel tryptic digestion as previously described (49).

### LC/MS/MS analysis

An aliquot of the tryptic digest (in 2 % acetonitrile/0.1% formic acid in water) was analyzed by LC/MS/MS using an Orbitrap Fusion™ Tribrid™ mass spectrometer (Thermo Scientific™) interfaced with a Dionex UltiMate 3000 Binary RSLCnano System. Peptides were separated with a C18 reversed-phase column (100 μm ID x 25 cm, 5 μm / 18Å Reprosil-Pur C18-AQ beads from Dr Maisch, Ammerbuch-Entringen, Germany) at a flow rate of 350 nl/min. Gradient conditions were: 3%-22% B for 40 min, 22%-35% B for 10min, 35%-90% B for 10 min, followed by 90% B for 10 min (solvent A, 0.1 % formic acid in water; solvent B, 0.1% formic acid in acetonitrile). The peptides were analyzed using a data-dependent acquisition mode. The survey scan was performed with 120K resolution from 350 to 1500 m/z with an AGC target of 2e5 and a max injection time of 50 msec. The DDA cycle was limited to 3 seconds. Monoisotopic masses were then selected for further fragmentation for ions with 2 to 5 plus charge within a dynamic exclusion range of 20 seconds. Fragmentation priority was given to the most intense ions. Precursor ions were isolated using the quadrupole with an isolation window of 1.6 m/z. HCD was applied with a normalized collision energy of 34%, and resulting fragments were detected using the rapid scan rate in the ion trap. The AGC target for MS/MS was set to 1e4, and the maximum injection time was limited to 35 msec.

The raw data files were processed using Thermo Scientific™ Proteome Discoverer™ software. The spectra were searched against the Uniprot-Homo sapiens database using Sequest. The database search was restricted to the following parameters. Trypsin was set as the enzyme with maximum missed cleavages set to 2. The precursor ion tolerance was set to 10 ppm, and the fragment ion tolerance was set to 0.8 Da. Carbamidomethylation on cysteine was selected as a static modification. Variable modifications were set to oxidation of methionine, acetylation of N-terminus, ubiquitination of Lysine (Gly-Gly), and SUMOylation of Lysine (T-Gly-Gly). The search results were validated and trimmed to a 1% FDR for strict conditions and 5% FDR for relaxed conditions using Percolator.

### Monitoring cellular SUMOylation of EPAC1 by SUMO2/3 affinity pull-down

C-terminally EYFP-tagged EPAC1 WT, K561R, or APEX-2 tagged EPAC1 WT or SIM/5A was transfected into HEK293 cells, respectively. Forty-eight hours after transfection, cells with 80-90% confluence were subject to heat shock treatment at 43 °C for 30 min, while control cells were kept at 37 °C. For each condition, two 10 cm plates were used. The cells were rinsed with cold PBS and lysed in lysis buffer containing 50 mM tris (pH 7.5), 150 mM NaCl, 1.5 mM MgCl_2_, 0.5 mM EDTA, 20 mM NEM, 1 mM phenylmethylsulfonyl fluoride (PMSF), 1% NP-40, and cOmplete Protease Inhibitor Cocktail on ice for 5 to 10 min. Cell lysates were harvested by centrifugation at 16,000g for 15 min to remove cell debris. Cell lysates with an equal amount of total cellular proteins (1.6 mg) were incubated with 3 μg of anti-SUMO2/3 antibody (Catalog no. M114-3, MBL Life Science) or mouse immunoglobulin G (IgG) (Catalog no. sc-2025, Santa Cruz Biotechnology) with gentle mixing at 4°C for 2 hours. Protein A/G PLUS-Agarose beads (20 μl), equilibrated in lysis buffer, were added to the sample mixtures and incubated at 4 °C with gentle mixing for 1 hour. The agarose beads were collected by centrifugation at 1,000g for 3 min and washed five times with buffer containing 50 mM tris (pH 7.5), 150 mM NaCl, 1.5 mM MgCl_2_, 0.5 mM EDTA, 20 mM NEM, 1 mM PMSF, 0.75% NP-40, and 5% glycerol. After the final wash, the beads were resuspended in 40 μl of 2× SDS sample buffer. The SUMO2/3 immunoprecipitation samples were analyzed by SDS-PAGE, followed by immunoblotting using anti-EPAC1 antibody 5D3 (Catalog no. 4155, Cell Signaling Technology).

### Confocal fluorescence microscopic imaging and analysis

HEK293 cells were plated on glass coverslips coated with 2% gelatin or poly-l-lysine (10 μg/ml) and transfected with EPAC1-EYFP and pcDNA-mRuby-UBA2. Twenty-four hours post-transfection, the cells were subjected to heat shock treatment at 43 °C for various time points, fixed with 4% PFA for 20 min at 37°C, rinsed three times with PBS for 5 min each, and stained with DAPI solution. Coverslips were mounted with FluorSave reagent for fluorescence microscopic imaging with a Nikon AXR or A1R laser confocal microscope system using the same parameter settings, including the laser power and exposure time. Images for more than eight randomly selected fields from at least three independent coverslips per treatment condition were collected.

### Intracellular cAMP Determination

The intracellular cAMP of HUVEC or HEK-293 cells with or without heat shock treatment was measured using a Direct cAMP ELISA kit from Enzo Biochem (Catalog no. AD1-900-066A) following the manufacturer’s instructions. The relative cAMP level was normalized to protein concentration.

### Cellular Rap-GTP pull-down assay

The cellular activities of Rap1 and Rap2 in HUVEC or HEK293 cells ectopically expressing EPAC1-EYFP, EPAC1(K561R)-YFP or EPAC1(K4R)-YFP were assessed using a glutathione S-transferase fusion of the Rap1-binding domain of RalGDS as described earlier (27). Briefly, cells were grown to 75% confluence in 10-cm Corning culture dish and subjected to heat shock treatment at 43 °C for 30 min while control cells were kept at 37 °C. Following three washes in PBS, the cells were lysed in a buffer containing 50 mM tris (pH 7.5), 150 mM NaCl, 2.5 mM MgCl_2_, 0.5% Na-deoxycholate, 0.1% SDS, 20 mM NEM, 1 mM PMSF, 1% NP-40, and 1 × Roche EDTA-free protease inhibitors. The cell lysate was mixed with 30 μl of glutathione-Sepharose beads with 30 μg of glutathione S-transferase-RalGDS-Rap1-binding domain bound and incubated at 4 °C for 2 h with gentle agitation. Following five washes in lysis buffer, the beads were suspended in 40 μl of SDS sample buffer. 10-15 μl of protein samples were loaded onto a 15% SDS-polyacrylamide gel and further analyzed with Western blot using Rap1 and Rap2 specific antibodies.

### Molecular modeling and dynamics simulation analyses of K561-SUMOylated EPAC1

The Rap1 bound EPAC1 structure was based on a published EPAC1 model based on homology modeling and rigid-body and ensemble analyses of Small-Angle X-ray Scattering (SAXS) data (50). In brief, the EPAC2:cAMP:Rap1 3CF6 PDB structure was used as the template for generating an EPAC1:cAMP:Rap1 model using Modeller (51). The SUMO molecule from the 3UIP structure was extracted and positioned near K561 of EPAC1 using Pymol. The linkage between the SUMO di-glycine terminus and EPAC1 K561 was made in COOT. The SUMO:EPAC1 interface residues were energy minimized with stereochemical restraints in COOT (52). This starting model was then prepared for NAMD (53) using the VMD solvation and autoionization modules (54) using a 30 Å padded box with a 150 mM NaCl concentration. The solvated model was then subjected to minimization and annealing in NAMD before performing a full Molecular Dynamics Simulation run of 30 ns. These were performed using an NPT ensemble and PME electrostatics with a 12 Å cutoff and 10 Å switching distance. We performed three MD Simulations to probe the SUMOylated EPAC1 Rap1 effector binding system. For the EPAC1-SUMO:Rap1 model. The choice of an initial position of the SUMO in the EPAC1:cAMP:Rap1 receptor-bound complex is highly restricted due to steric hindrance. The SUMO was manually positioned in a cleft between the Rap1 and RA domain of EPAC1 in the only possible orientation, one in which the EPAC1 N-terminal regulatory domains were distal to the SUMO and could not interact. Therefore, the MD Simulations were performed with the EPAC1 C-terminal domains (EPAC1^CT^) that span K561 by truncating the N-terminal regulatory domains. For the EPAC1^CT^-SUMO model, the initial position of the SUMO relative to the EPAC1^CT^ domains is not restrained by its covalent bond to K561 of EPAC1. The EPAC1^CT^-SUMO model was derived from a minimized and annealed EPAC1^CT^-SUMO:Rap1 model with the Rap1 effector removed. This gave the SUMO an initial orientation to the EPAC1^CT^ domains, which was physically reasonable. The MD Simulation run of this EPAC1^CT^-SUMO model probes whether this was a stable orientation that may be adopted before receptor binding or if the SUMO would move from this position. The third simulation was performed on the full-length ternary EPAC1:cAMP:Rap1 homology model reported previously (50). The resulting ternary model was used for comparative analysis with the SUMOylated model. This full-length ternary model after MD improved fitting to the experimental SAXS data, validating the effectiveness of the MD analyses.

### Statistical Analyses

Results are presented as mean ± standard error of the mean (SEM). Data was analyzed for normality and equal variance using the Shapiro-Wilk normality test and an F-test, respectively. For data exhibiting normal distribution, a Student t-test was implemented to compare two groups of equal variances, whereas a Welch t-test was used in cases with unequal variance. For non-normal distributions, a Mann-Whitney test was conducted to compare groups. Additionally, one-way ANOVA with a Bonferroni post hoc test was used to compare groups of three or more with normal distributions. A p-value less than 0.05 was considered as statistically significant.

## Acknowledgments

Grants from the National Institutes of Health R35GM122536 and from the American Heart Association 20TPA35410051 supported this work. The authors thank the Clinical and Translational Proteomics Service Center at the University of Texas Health Science Center at Houston for the proteomic analysis and Dr. Olga Chumakova and Ms. Zhengmei Mao from UTHealth Center for Advanced Microscopy for technical assistant. The authors acknowledge the Sealy Center for Structural Biology and Molecular Biophysics at the University of Texas Medical Branch at Galveston for providing research resources. The funders had no role in the study design, data collection and analysis, decision to publish, or manuscript preparation.

## Author Contributions

Wenli Yang: Methodology; Investigation; Validation; Formal analysis; Data Curation; Writing-Method, Review & Editing; Visualization. Fang Mei: Methodology; Investigation; Validation; Formal analysis; Data Curation; Writing-Review & Editing; Visualization. Wei Lin: Methodology; Investigation; Validation; Data Curation; Writing-Method. Mark White: Methodology; Investigation; Validation; Formal analysis; Data Curation; Writing-Method, Review & Editing; Visualization. Li Li: Methodology; Investigation; Validation; Formal analysis; Data Curation; Writing-Method, Review & Editing. Yue Li: Methodology; Resources. Sheng Pan: Data Curation; Writing-Method, Review & Editing. Xiaodong Cheng: Conceptualization; Methodology; Investigation; Validation; Formal analysis; Resources; Data Curation; Writing-Original draft preparation, Review & Editing; Visualization; Supervision; Project administration; Funding acquisition.

## Competing Interest Statement

The authors declare no competing financial interests.

## Notes

### Competing Interest Statement

The authors have declared no competing interest.

