## Supplemental Data for "Protein SUMOylation promotes cAMP-independent EPAC1 activation"

[#]These authors contributed equally.

[§]Current Address: Cell Therapy Manufacturing Center. 2130 W Holcombe Blvd, Houston TX, 77030.

*Corresponding author: Xiaodong Cheng.

**Supplementary Data**

**Table S1. Predicted SUMOylation sites and SUMO-interacting motifs.**

| Putative SUMOylation site | | |
|---|---|---|
| **Position** | **Sequence** | **Database Hit** |
| K212 | KAVAHLSNSVKRELAAVLLFE | 13 |
| K310 | FNRIIKDVEAKTMRLEEHGKV | 1 |
| K319 | AKTMRLEEHGKVVLVLERASQ | 1 |
| K517 | GSSCAIQVGDKVPYDICRPDH | 3 |
| K561 | DGWTKGQVLVKVNSAGDAIGL | 1 |
| K698 | PRAQLLRKFIKLAAHLKEQKN | 2 |
| K764 | NHRVYRLALAKLSPPVIPFMP | 1 |
| K778 | PVIPFMPLLLKDMTFIHEGNH | 1 |
| K864 | ASTWAYVQQLKVIDNQRELSR | 1 |
| **Putative SUMO-interacting motif** | | |
| **Position** | **Sequence** | **Score/Database Hit** |
| AA 320-323 | MRLEEHGKVVLVLERASQGA | 2.020/2 |
| AA 321-324 | RLEEHGKVVLVLERASQGAG | 3.166/0 |

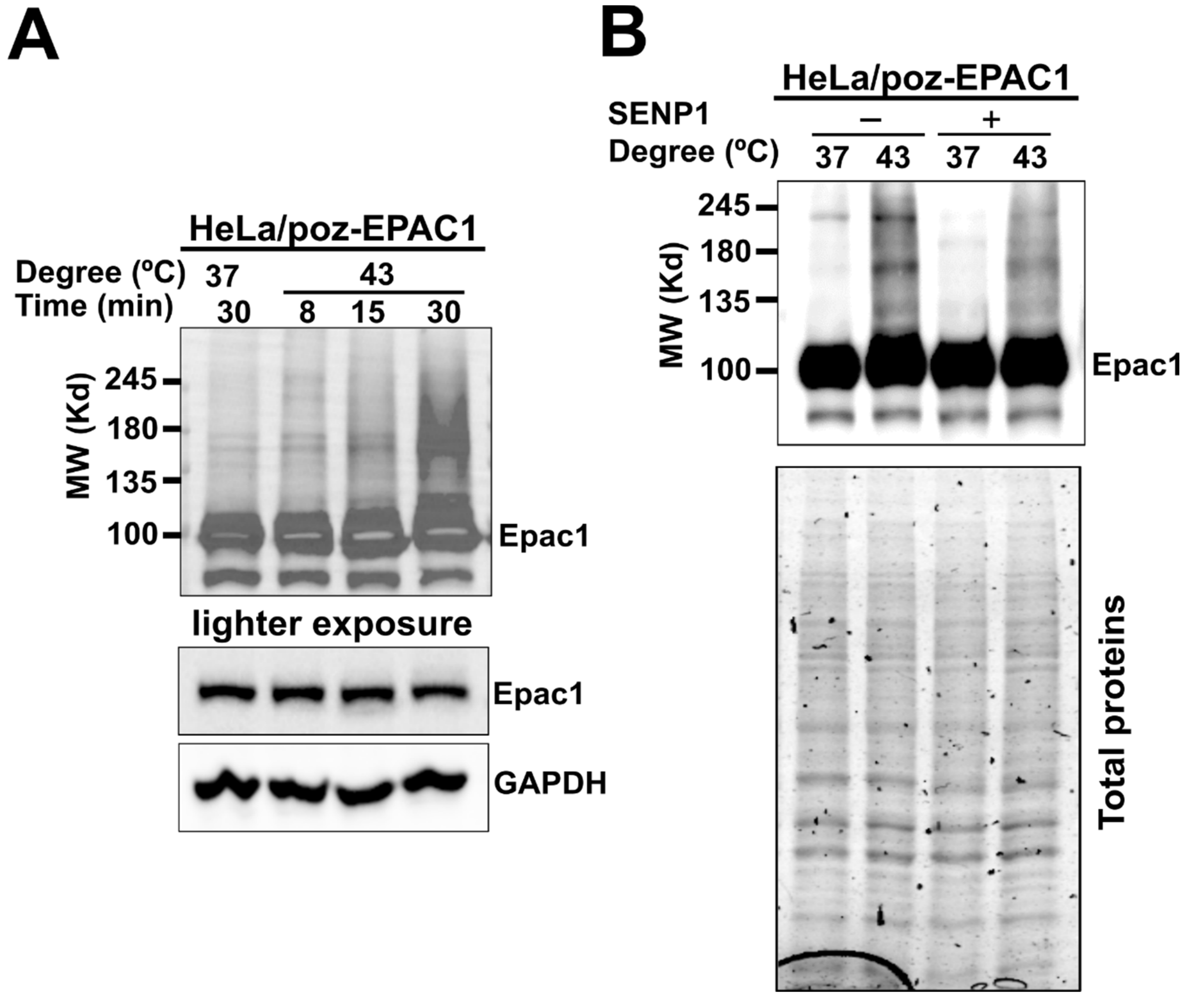


**Figure S1. Heat shock promotes EPAC1 post-translational modifications (PTM).** (A) Levels of cellular EPAC1 PTM probed by immunoblotting using anti-EPAC1 antibody in HeLa/poz-EPAC1 in response to heat shock as a function of time. (B) Levels of cellular EPAC1 PTM in HeLa/poz-EPAC1 cells probed by immunoblotting using anti-EPAC1 antibody, with and without heat shock (30 min) and with or without SENP1 (220 nM) treatment at 37 °C for 20 min. Similar results were obtained from at least three independent experiments.

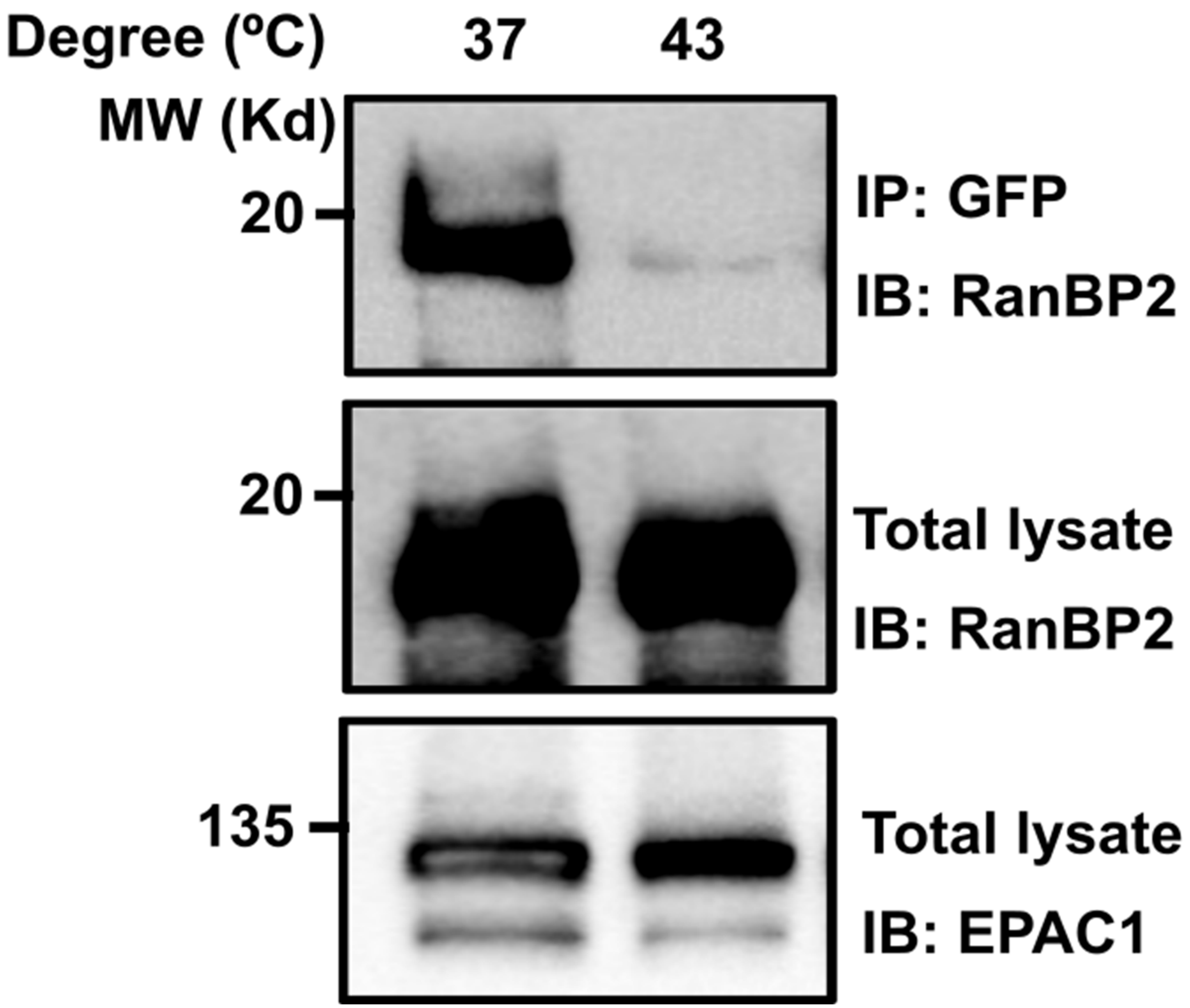


**Figure S2. Heat shock reduces EPAC1 RanBP2 interaction.** Interaction between ectopically expressed EPAC1-EYFP and endogenous RanBP2 in response to heat shock as probed by affinity purification using anti-GFP antibodies.

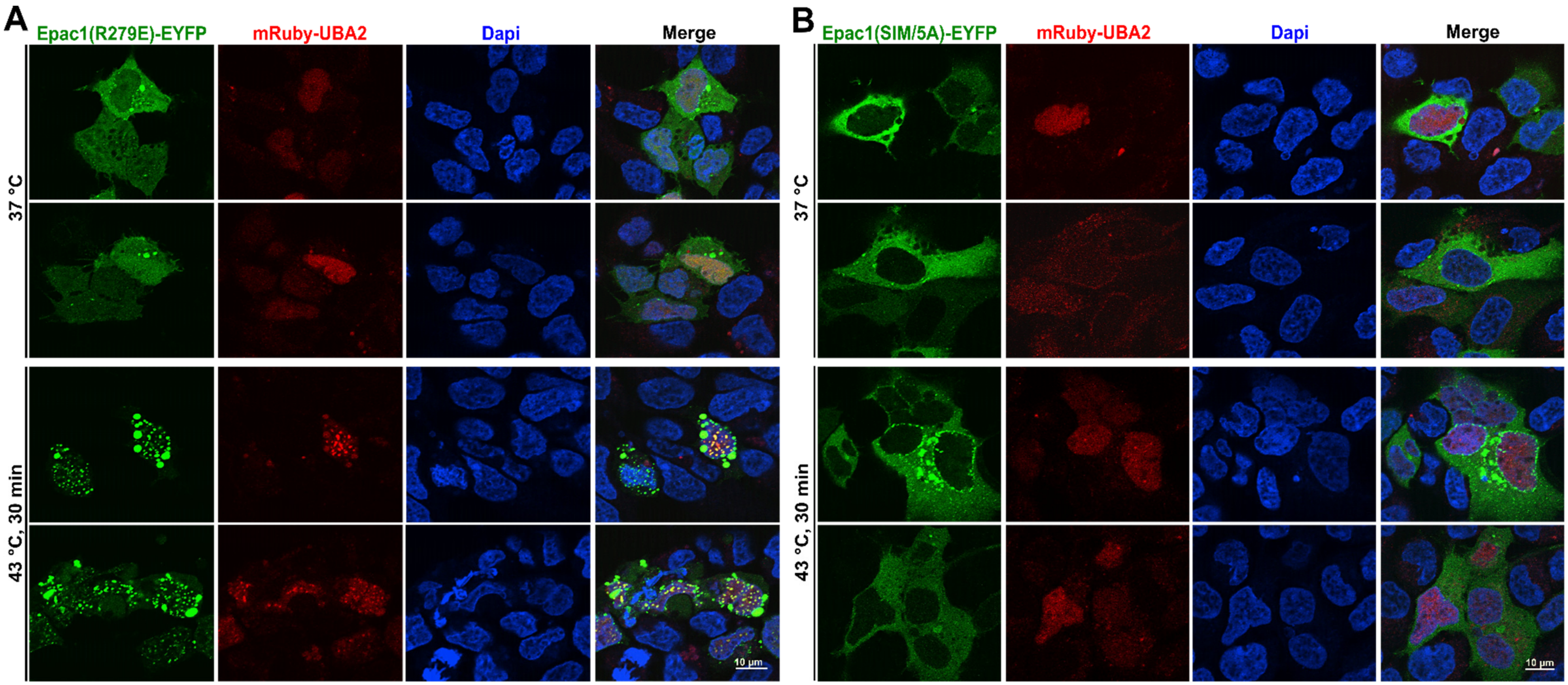


**Figure S3. Changes in cellular distributions of EPAC1 mutants in response to heat shock.** Confocal images of HEK293 cells ectopically expressing EPAC1(R279E)-EYFP (A) or EPAC1(SIM/5A)-EYFP (B) in response to heat shock treatment at 43 °C for 30 min.

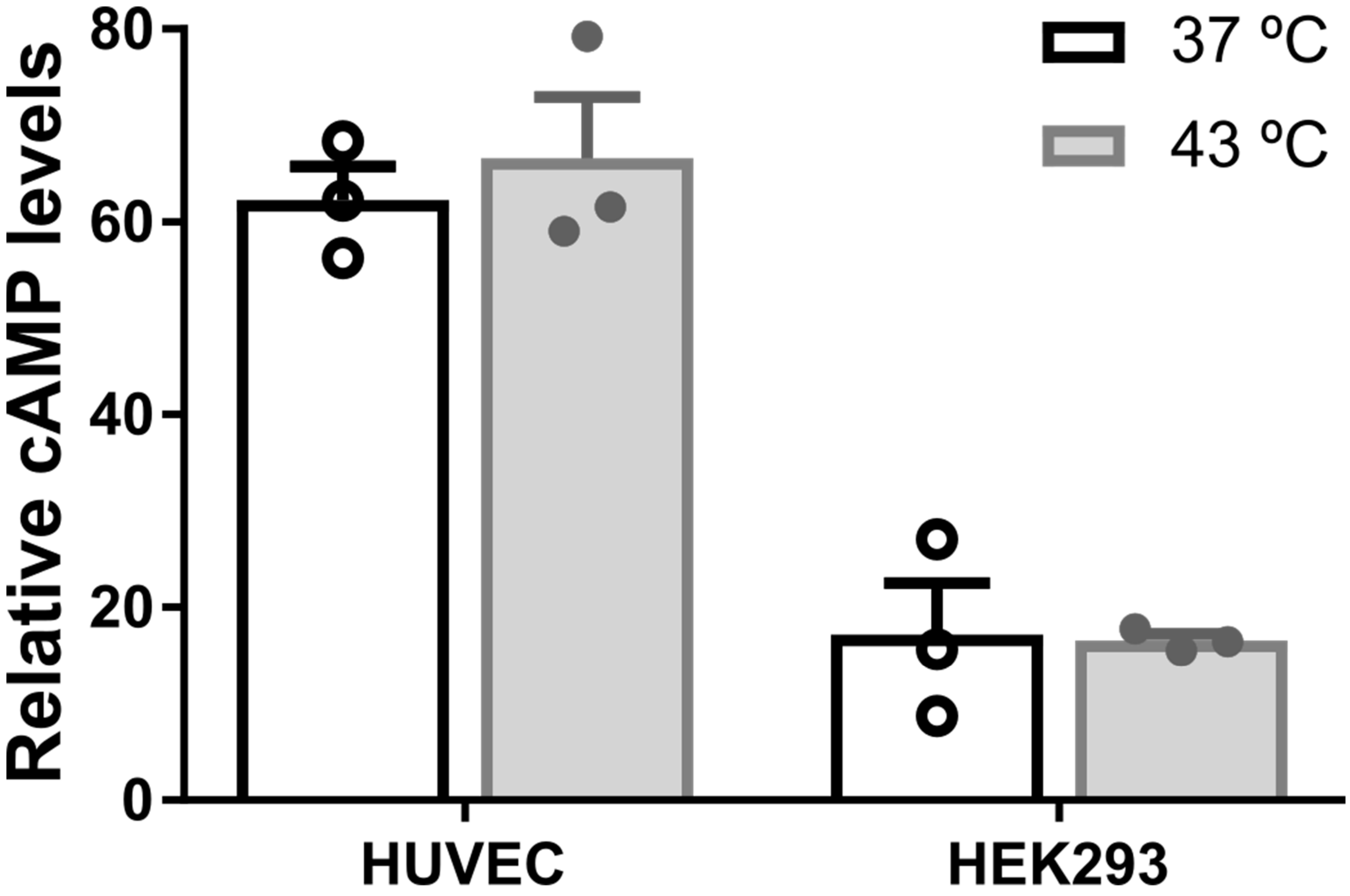


**Figure S4. Effects of heat shock on levels of intracellular cAMP.** Levels of cellular cAMP in HUVEC and HEK293 cells at 37 °C or after heat shock at 43 °C for 30 min.

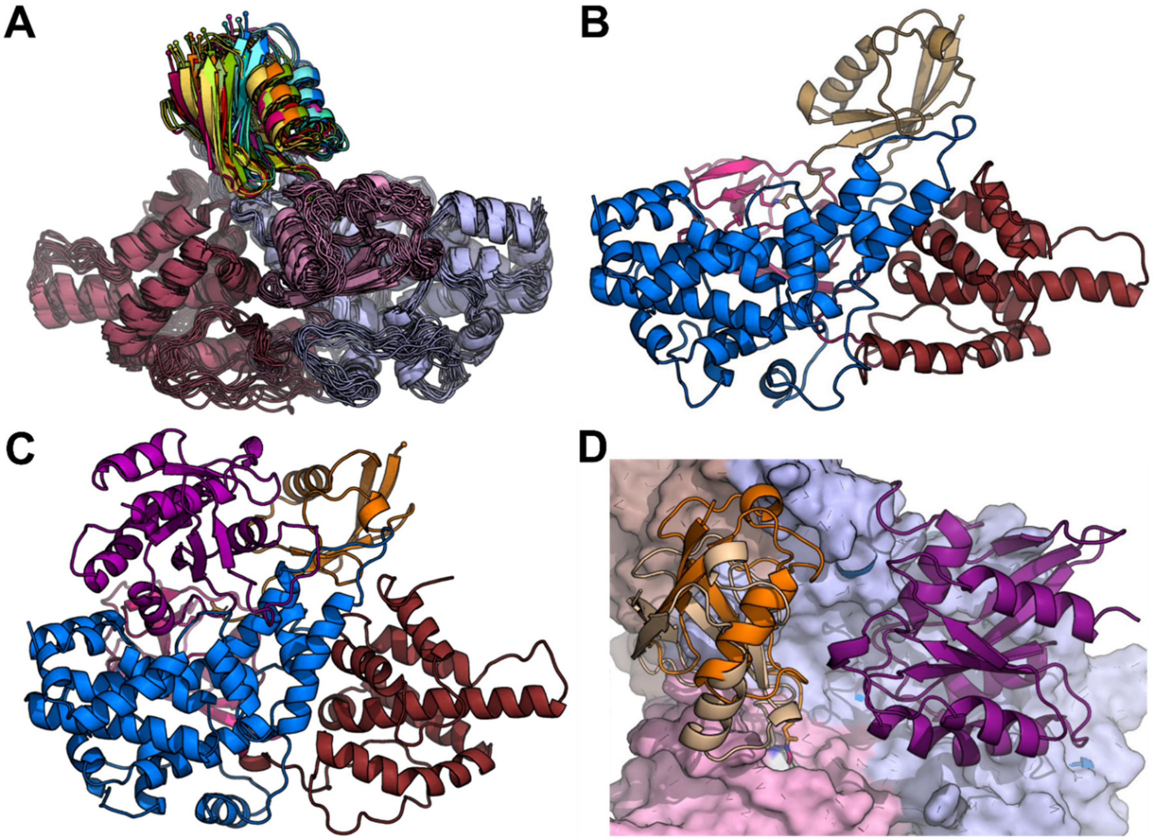


**Figure S5. The EPAC1$^{CT}$-SUMO:Rap1 and EPAC1$^{CT}$-SUMO models after 30ns of Molecular Dynamics.** (A) SUMO orientations EPAC1$^{CT}$-SUMO during the 30 ns MD. The EPAC1$^{CT}$-SUMO MD model is shown colored by domain: REM: brown, RAP: pink, and GEF: blue. The SUMO is heat map colored by MD time from blue (0) to red (30 ns). (B) The final EPAC1$^{CT}$-SUMO MD model, with the SUMO colored in beige. (C) The final EPAC1$^{CT}$-SUMO:Rap1 MD model, with the SUMO colored in orange and Rap1 colored in purple. (D) SUMO orientation shifts slightly in the absence of the Rap1 domain, but it remains in roughly the same position. The EPAC1$^{CT}$-SUMO MD model is shown in light tones with the SUMO colored in beige and EPAC1 surfaced, while the EPAC1$^{CT}$-SUMO:Rap1 MD model is in bold tones with the SUMO colored in orange and Rap1 colored in purple.

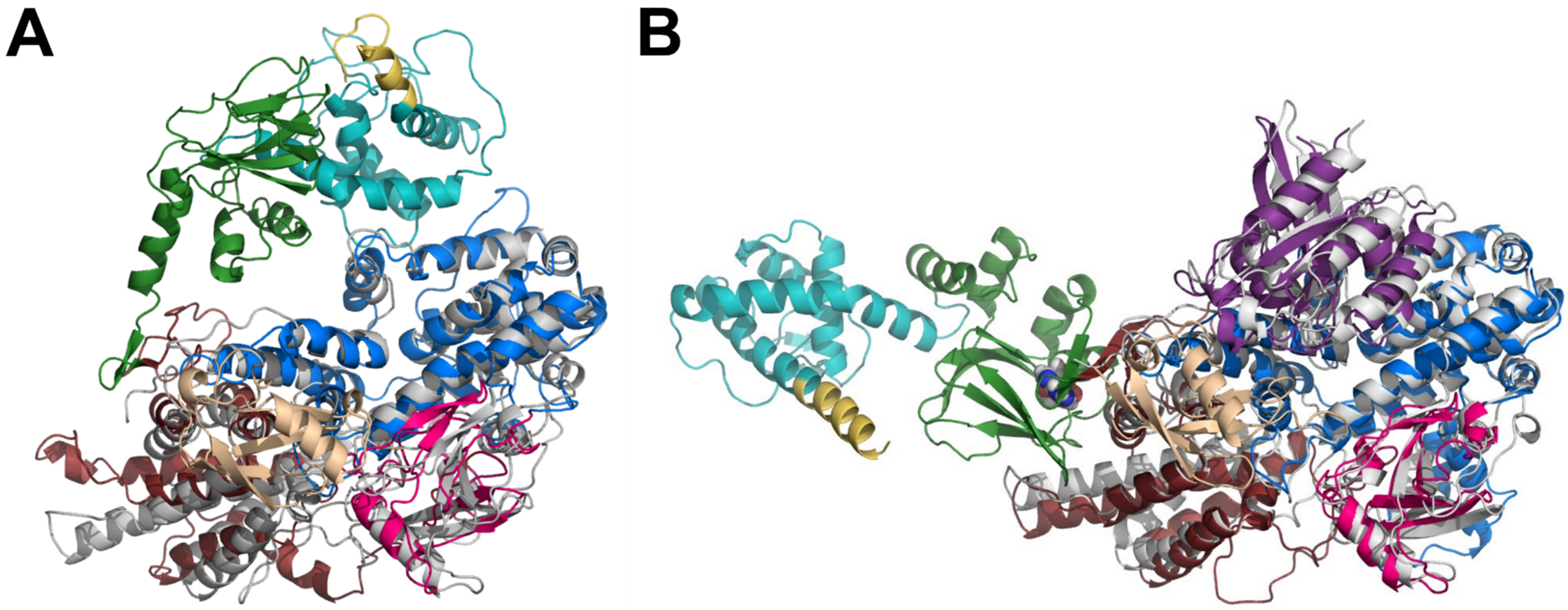


**Figure S6. The MD models of EPAC1$^{CT}$-SUMO and EPAC1$^{CT}$-SUMO:Rap1.** (A) The EPAC1$^{CT}$-SUMO model with EPAC$^{CT}$ colored in grey and SUMO colored in beige superimposed with the apo-EPAC1 model (DEP: cyan; CNBD: green; REM: maroon; RA: pink; CDC25HD: blue). (B) The EPAC1$^{CT}$-SUMO:Rap1 model with EPAC$^{CT}$ colored in grey, SUMO colored in beige, and Rap1 colored in purple, superimposed with the ternary EPAC1:cAMP:Rap1 model.
